# Na^+^/K^+^-ATPase Blockade Reversibly Attenuates Proliferation and Metabolic Function in K562 Human Myelogenous Leukemia Cells

**DOI:** 10.64898/2026.09.11.751086

**Authors:** Simon R Bowden, Eli T Colton, Amber D Norgaard, McKenzie M Haas, Erynn R Dunnigan, Carly J Goven, Grace O Goven, Thomas J Hinnenkamp, Keagan D Walker, Theresa M Ballance, Blaise I Kathol, Elizabeth R Murphy, Kristian M Van Slyke, Rylie J Webb, Ashlin B Schaefbauer, Emily R Biggane, Joseph P Biggane

## Abstract

The Na^+^/K^+^-ATPase consumes a substantial portion of cellular ATP to maintain essential electrochemical gradients, yet its role in regulating cancer cell proliferation and phenotypic plasticity remains complicated by its dual function as an ion transporter and a scaffolding receptor. Here, we investigate the physiological consequences of pharmacological NKA blockade using digoxin in K562 human chronic myelogenous leukemia cells. Submicromolar digoxin exposure induced a concentration-dependent attenuation of cell proliferation (IC_50_= 151.0 nM) and a statistically significant depression of metabolic reducing capacity without impairing cell viability. Competitive supplementation with extracellular potassium salts produced a surmountable rightward shift in digoxin sensitivity, confirming that growth inhibition is driven by on-target NKA pump occupancy rather than non-specific interactions. Furthermore, this cytostatic growth attenuation and metabolic depression were found to be fully reversible upon drug clearance. K562 cells are largely refractory to canonical kinase-overdrive stress checkpoints as they harbor constitutive BCR-ABL tyrosine kinase activity alongside non-functional TP53 and a homozygous deletion of CDKN2A (p16^INK4a^). Consequently, these findings suggest that NKA perturbation attenuates K562 growth primarily through ion dyshomeostasis and secondary active transport constraints rather than scaffold-mediated signaling. Overall, we present a novel model for elucidating the role of the NKA in cancer proliferation and dissecting the mechanistic underpinnings of ion-mediated alterations in cancer cell physiology.

## Introduction

Even as cancer therapies gain precision and efficacy, the rapid diversification of tumor populations often leads to therapeutic resistance, especially in recurrent populations where resistance has been actively selected [1,2]. Thus, there is a growing interest in the promise of identifying therapeutic avenues that constrain phenotypic plasticity in cancer cells, yet actionable therapeutic levers remain elusive [3]. Ion transport is often highly dysregulated in cancer and presents an exploitable vulnerability to manipulate the simultaneously elevated energic demands and biosynthesis requirements necessary for uncontrolled, aberrant cellular proliferation [4].

The molecular machinery for ion handling represents an estimated 5% of all protein-coding genes in the human genome, and maintaining functional transmembrane ion gradients comprises approximately one-third of the total ATP consumption of a typical mammalian cell [5,6]. Ion transport gene expression dysregulation has been demonstrated across the cancer landscape with implications as a key driver of many cancer hallmarks, and even suggestions that certain cancers are so-called “oncochannelopathies” [1,4]. Further, intracellular ion concentration, ion flux, and membrane potential are associated with cell cycle progression in both normal cells and in cancer [7,8]. The exact intracellular mechanisms underlying ion-mediated changes in cancer cell physiology and function remain elusive, yet precise control of intracellular ion balance drives maintenance of cellular homeostasis and several critical cell functions, such as: protein stability and function, cell volume and osmotic regulation, secondary active transport of nutrients and waste, and cell signaling [9,10]. While hundreds of different ligand- and voltage-gated channels, ion pumps, and transporters facilitate movement of ions down concentration gradients to drive energetically unfavorable reactions and conformations, the Na^+^/K^+^-ATPase (NKA) is a central cog maintaining precise intracellular sodium and potassium homeostasis while consuming at least one-fifth of a typical cell’s ATP [6].

Despite substantial toxicity challenges associated with targeting the ubiquitous NKA, perturbing its activity remains a powerful approach for dissecting how intracellular ion dynamics dictate cancer cell physiology [11,12]. Elucidating downstream ion-mediated effects in cancer is further complicated by the NKA’s dual role as an ion transporter and scaffolding receptor [13,14]. In particular, the signalosome hypothesis, made popular by Xie and colleagues, posits that the obligatory α subunit of the NKA heteromeric protein complex binds to the proto-oncogene tyrosine kinase, Src [15,16]. In this model, cardiac glycoside binding to NKA induces a conformational change that releases activated Src [17]. Mechanistically, the signalosome hypothesis posits that ligand-induced Src activation triggers EGFR transactivation, downstream MAPK and PI3K/Akt signaling cascades, and reactive oxygen species generation, which together drive cytostatic growth arrest or apoptotic pathways [14,18]. Thus, while it has been demonstrated in several cancer cell types that NKA manipulation can decrease cancer cell proliferation and viability, it remains difficult to disentangle Src-mediated effects from ion transport-mediated effects [19-24].

The K562 human chronic myelogenous leukemia cell line presents a distinct opportunity to study the role of the NKA in cancer cells. K562 cells are positive for the BCR-ABL fusion mutation, conferring constitutive tyrosine kinase expression and activity that drives autonomous proliferation and suppresses canonical apoptotic responses to growth-factor withdrawal and cellular stress [25-27]. Further, K562 cells carry an inactivating mutation in TP53 and homozygous deletion of the CDKN2A locus (p16^INK4a^), disabling canonical checkpoint and growth-arrest machinery [28,29]. Taken together, any anti-proliferative response observed in K562 cells following NKA inhibition can be attributed primarily to genuine biophysical ion dyshomeostasis or to non-canonical, receptor-mediated stress signaling via the NKA–Src signalosome.

In this study, we investigated the relationship between cancer cell ion dyshomeostasis and cancer cell proliferation. To alter ion homeostasis in cells, we targeted the NKA in K562 cells using the cardiac glycoside digoxin. We hypothesized that NKA blockade would result in a reversible decrease in K562 cell proliferation. Our experiments focused on K562 proliferation, viability, and metabolic capacity in response to a wide range of digoxin concentrations.

Additionally, we confirmed that the observed effects of digoxin were NKA-dependent and assessed the reversibility of these cellular phenotypes. Our results demonstrate that K562 cells are a powerful model for elucidating the role of the NKA in cancer proliferation and disentangling the mechanistic underpinnings of ion-mediated alterations in cancer cell physiology.

## Materials and Methods

### Reagents

Iscove’s Modified Dulbecco’s Medium, Hyclone Fetalclone III, and Antibiotic/Antimycotic Solution were obtained from Cytiva. Pure molecular grade ethanol and potassium chloride were obtained from Fisher Scientific. Potassium gluconate and sodium gluconate were obtained from TCI. Dulbecco’s phosphate buffered saline was obtained from Corning. 0.4% trypan blue solution was obtained from Gibco. Resazurin (Alamar Blue) ready-to-use solution was obtained from ApexBio Technology. Digoxin was obtained from Acros Organics.

### Cell Culture

Human chronic myelogenous leukemia cells (K562) were obtained from Sigma-Aldrich and maintained at 37°C/5% CO_2_ in a CO_2_ incubator (Shel Labs). Cells were maintained in Iscove’s modified Dulbecco’s medium, with 10% synthetic Hyclone Fetalclone III and Antibiotic-Antimycotic solution. All cells used in these experiments were from P3-P21.

### Cell Growth and Viability Experiments

K562 cells were harvested at log-phase concentrations, then plated in triplicate wells at 30,000 cells/mL in complete media containing 80% ethanol vehicle or 0.1 nM, 1 nM, 10 nM, 100 nM, 1 μM, or 10 μM digoxin. All experiments using digoxin were diluted from a stock solution prepared in 80% molecular-grade ethanol. Serial dilutions were prepared to ensure each concentration was administered in 80% ethanol vehicle and <1% v/v bolus. Experiments using 20 mM potassium gluconate, 20 mM sodium gluconate, or 20 mM potassium chloride were completed by dissolving each salt into complete media. Cells were incubated over 96-hours, then each triplicate was counted using a TC-20 automated cell counter (Bio-Rad) with trypan blue to measure viability. Lower concentrations (0.1-10 nM) produced no detectable changes in viability relative to vehicle, so data were omitted from bar plots for clarity. Triplicate cell counts and viability were averaged as technical replicates. Biological replicates represent independent experiments conducted from separate culture flasks maintained across distinct passage generations.

### Recovery Assay

K562 cells were harvested following a 96-hour exposure to either 80% ethanol vehicle or 100 nM Digoxin at an initial seeding density of 100,000 cells/mL. A higher density of cells was used for this set of experiments due to an uncertainty as to what degree 100 nM digoxin would inhibit cell growth, and to ensure a quantifiable cell population. Upon completion of exposure, cells were harvested by centrifugation, washed with PBS, resuspended with fresh complete culturing media, and plated in triplicate wells at 100,000 cells/mL. Following 96-hours of recovery in fresh media, each triplicate was counted using a TC-20 automated cell counter with trypan blue to measure viability. Triplicate cell counts and viability were averaged as technical replicates.

Biological replicates represent independent experiments conducted from separate culture flasks maintained across distinct passage generations.

### Metabolic Reducing Capacity Assay

Initial set up for these experiments was identical to the cell growth and viability experiments. Briefly, K562 cells were harvested at log-phase concentrations then plated in triplicate wells at 30,000 cells/mL in complete media containing 80% ethanol vehicle or 0.1 nM, 1 nM, 10 nM, 100 nM, 1 μM, or 10 μM. All experiments using digoxin were diluted from a stock solution prepared in 80% molecular-grade ethanol. Serial dilutions were prepared to ensure each concentration was administered the same bolus of 80% ethanol vehicle. Cells were incubated over 96-hours, harvested, resuspended in complete culturing media, then counted using a TC-20 automated cell counter (Bio-Rad) with trypan blue to measure viability. Cells were plated in triplicate at 50,000 cells/well per manufacturer recommendation because of low cell growth at high digoxin concentrations, then incubated with resazurin (alamar blue) overnight. Endpoint absorbance was measured at 570 nm and 600 nm on a multimode microplate spectrophotometer (Accuris Instruments). Metabolic reduction was quantified as the A_570_/A_600_ absorbance ratio subtracted by the cell-free media blank ratio, then expressed as a percentage relative to the matched vehicle control. Lower digoxin concentrations (0.1-10 nM) produced no detectable changes in metabolic reducing capacity relative to vehicle, thus data were omitted from bar plots for clarity. Biological replicates represent independent experiments conducted from separate culture flasks maintained across distinct passage generations.

### Statistical Methods

For all dose-response assays, internal technical triplicates from each multi-well plate were averaged to calculate a single value per independent biological replicate, and these values were pooled to determine the mean ± SEM across biological runs. Dose-response curves were generated in Prism 11 (GraphPad) using a 3-variable non-linear regression analysis. Dose-response curves were calculated after being normalized to vehicle-only control cell counts. Since a null concentration cannot be plotted on a logarithmic plot, the top of each dose-response curve was constrained to 100%. IC50 and maximum efficacy parameters were derived from the generated non-linear regression line equation. All graphical representations of individual data points are presented as mean ± SEM. Values derived from the line of the non-linear regression are presented as the mean value, 95% CI, and R^2^.

Discrete viability, metabolic, and recovery assays are presented as mean ± SD in the text and shown as mean ± SEM in graphical representations. Values were internally normalized to matched vehicle-only controls within each independent experiment. Vehicle controls are displayed at 100% without error bars to establish the visual reference baseline. Statistical comparisons between groups were assessed using ordinary 1-way ANOVA and Dunnett’s multiple comparisons test. Statistical significance was determined using the following α thresholds P< 0.05 (*), P< 0.01 (**), P< 0.001 (***), P< 0.0001 (****). All graphical representations of these data are presented as the mean ± SEM.

## Results

### Digoxin Exposure Induces Concentration-Dependent Growth Inhibition in K562 Cells

To elucidate how ion manipulation influences cancer cell growth, K562 cells were plated at 30,000 cells/mL and exposed to 0.1 nM-10 μM digoxin. 96-hour digoxin exposure resulted in a dose-dependent decrease in growth of K562 cells, with 1 μM and 10 μM digoxin exhibiting no substantial growth (Figure 1A). Non-linear regression analysis of end-point cell growth revealed a maximal growth inhibition of 89.52% (95% CI= 84.00-95.19%, R^2^=0.9966) and IC50 concentration was determined to be 151.0 nM (95% CI= 110.0-205.2 nM, R^2^=0.9966) (Fig. 1B). We observed 98.22±1.42% viability relative to vehicle-only control at 100 nM digoxin, while 1 μM digoxin resulted in 39.51±6.68% viability relative to vehicle-only control, and 10 μM digoxin resulted in 47.32±16.54% viability relative to vehicle-only control. Both 1 μM and 10 μM digoxin exposure resulted in statistically significant decreases in cell viability (Fig. 1C). Taken together, these data demonstrate a dose-dependent non-lethal reduction in K562 cell proliferation upon exposure to 100 nM digoxin.

**Figure 1.**
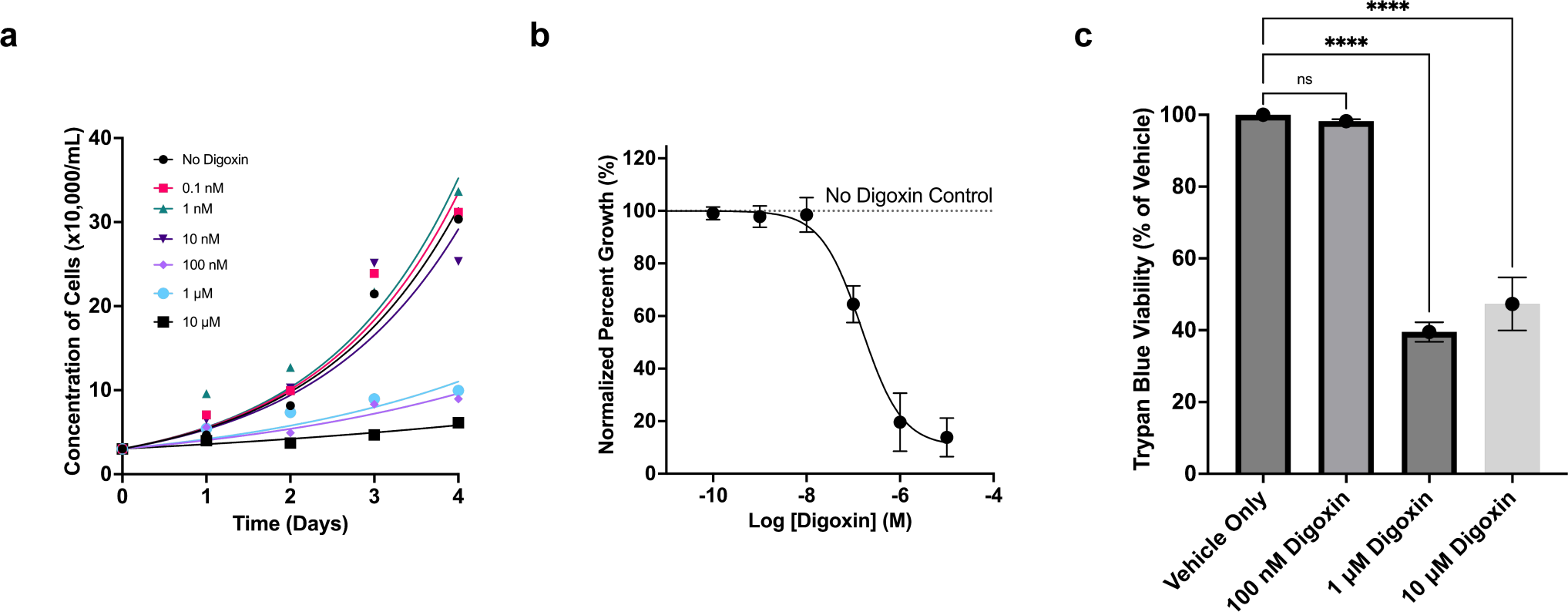
Digoxin-induced K562 growth inhibition and viability. (A) Representative growth curve of K562 cells from single biological replicates exposed to varying concentrations of digoxin. (B) A dose-response plot of K562 cell growth following 96-hour exposure to varying concentrations of digoxin. The y-axis values are normalized as a percentage of K562 growth with no digoxin exposure. All values are presented as the mean ± SEM. N=7 biological replicates, except N=6 for 1 nM and 1 μM digoxin and N=5 for 10 μM digoxin. Concentration-specific dropouts occurred due to contamination or unreadable counts. (C) Trypan blue indicated cell viability of selected K562 cell populations exposed to varying concentrations of digoxin for 96 hours. The number of biological replicates is identical to (B). All values are presented as mean ± SEM normalized to vehicle-only control growth. Statistical significance is represented as P< 0.05 (*), P< 0.01 (**), P< 0.001 (***), P< 0.0001 (****).

### Digoxin Target Specificity for NKA Indicated by Potassium-Selective Attenuation of Digoxin-Mediated Effects in K562 Cells

To better understand the mechanism of digoxin-mediated inhibition, we took advantage of the well-documented competitive inhibition of cardiac glycosides by elevated extracellular potassium [30,31]. Supplementation of K562 cells with 20 mM potassium gluconate, exposed to increasing concentrations of digoxin, resulted in a surmountable parallel rightward shift of the digoxin dose-response curve by two orders of magnitude (Fig. 2A). The IC50 of digoxin in the presence of 20 mM potassium gluconate was 1309 nM (95% CI=785.0-2219 nM, R^2^=0.9943).

**Figure 2.**
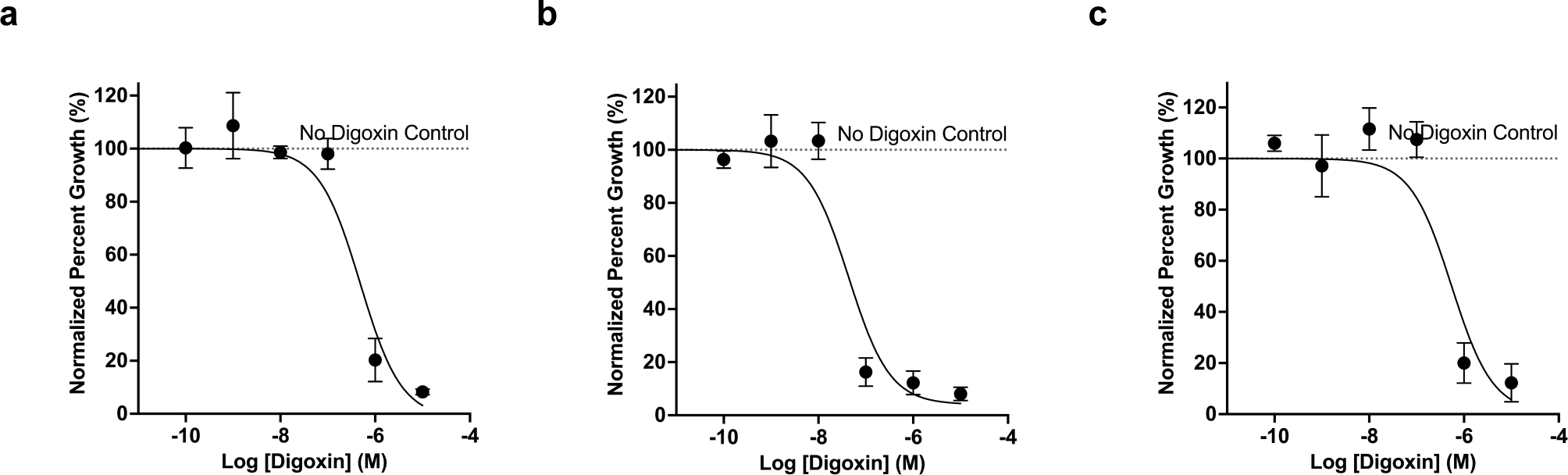
Ion competition and ion replacement demonstrate digoxin selectivity for NKA. (A) A dose-response plot of K562 cell growth following 96-hour exposure to varying concentrations of digoxin and 20 mM potassium gluconate. The y-axis values are normalized as a percentage of K562 growth with no digoxin exposure. All values are presented as the mean ± SEM. N=3 biological replicates for all concentrations. (B) A dose-response plot of K562 cell growth following 96-hour exposure to varying concentrations of digoxin and 20 mM sodium gluconate. The y-axis values are normalized as a percentage of K562 growth with no digoxin exposure. All values are presented as the mean ± SEM. N=3 biological replicates for all concentrations. (C) A dose-response plot of K562 cell growth following 96-hour exposure to varying concentrations of digoxin and 20 mM potassium chloride. The y-axis values are normalized as a percentage of K562 growth with no digoxin exposure. All values are presented as the mean ± SEM. N=3 biological replicates for all concentrations.

We did not observe the same digoxin desensitization in the presence of 20 mM sodium gluconate supplementation (Fig. 2B). The IC50 was calculated to be 43.36 nM (95% CI= 12.84-139.1 nM, R^2^= 0.9400). Supplementation with 20 mM potassium chloride resulted in an IC50 of 544.4 nM (95% CI= 121.2-2421 nM, R^2^= 0.9088) (Fig. 2C). Further, since gluconate is cell-impermeable and largely biologically inert, the coherence between the data generated in the presence of either potassium gluconate or potassium chloride supports the conclusion that potassium competes with digoxin to mediate the protective effect, rather than an osmotic phenomenon [9,10]. Collectively, these data strongly suggest that digoxin selectively acts upon NKA to inhibit K562 cell growth and potassium interferes with this inhibition.

### NKA Blockade Results in Attenuated Metabolic Reducing Capacity in K562 Cells

To investigate the potential mechanism(s) underlying the reduced proliferation observed when K562 cells are exposed to digoxin concentrations above 100 nM, we employed a resazurin assay to determine the cellular reducing capacity of these cells. Following a 96-hour exposure to vehicle or digoxin concentrations, 100 nM digoxin exposure resulted in 78.43±10.43% resazurin reduction, 1 μM digoxin exposure resulted in 14.07±10.09% resazurin reduction, and 10 μM digoxin exposure resulted in 11.83±7.32% resazurin reduction when normalized to the resazurin reduction of vehicle-only exposure (Fig. 3). All concentrations tested represented a statistically significant depression in resazurin reducing capacity when compared to vehicle-only control. Importantly, resazurin reduction is indicative of intrinsic cellular reducing capacity (NAD(P)H turnover), as each resazurin assay was seeded with equivalent live cell numbers regardless of the digoxin exposure concentration [32,33]. Taken together, these results suggest that digoxin exposure results in significant depression of metabolic reducing capacity in K562 cells independent of cell viability.

**Figure 3.**
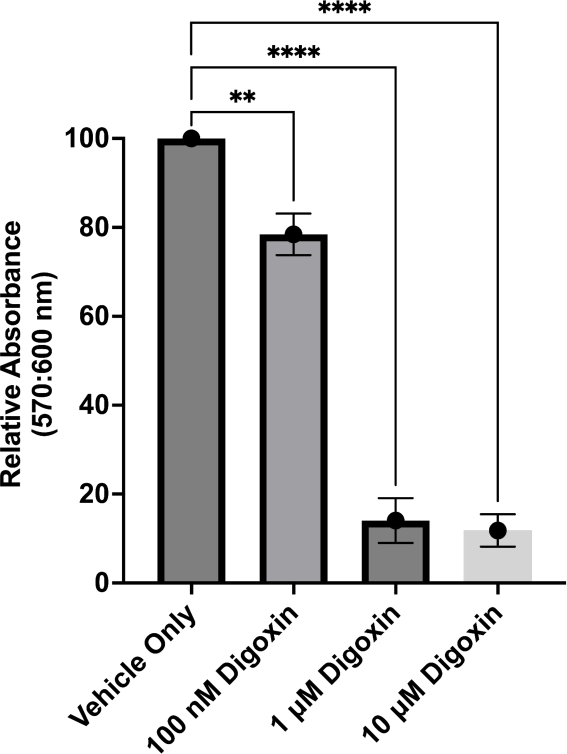
Metabolic reducing capacity of K562 cells exposed to digoxin. Plot showing resazurin reducing capacity of K562 cells exposed to selected concentrations of digoxin for 96 hours. Statistical significance is represented as P< 0.05 (*), P< 0.01 (**), P< 0.001 (***), P< 0.0001 (****). N=5 biological replicates for all concentrations, except N=4 for 1 μM and 10 μM digoxin. Concentration specific dropouts occurred due to unreadable counts.

### Growth and Metabolic Recovery of K562 Cells Reveals that the Effects of NKA Blockade are Transient

To further investigate whether significant damage is caused by digoxin exposure and the reversibility of digoxin-mediated growth and metabolic depression, we assessed the ability of K562 cells to continue growth once digoxin exposure ceased. Cells were subcultured after a 96-hour exposure to 100 nM digoxin or vehicle, then K562 cells were plated in fresh complete culture media, free of digoxin. To assess this, we compared growth of K562 cells exposed to 100 nM digoxin or vehicle after digoxin exposure and then again after an equivalent period of recovery (Fig. 4A). Following 96-hour exposure to 100nM digoxin, K562 cells exhibited 74.79±7.11% growth of vehicle-only controls. Following 96-hours recovering in fresh complete culture media, the same K562 cells exhibited 96.66±3.44% growth of vehicle-only control (Fig.4A). Thus, when compared, we observed a statistically significant recovery (P=0.019) in K562 cell growth to typical rates once digoxin was removed. When we assessed metabolic activity in recovered K562 cells, we observed 114.51±25.98% resazurin reducing capacity in cells exposed to 100 nM digoxin when compared to vehicle-only controls (Fig. 4B). This difference was statistically insignificant (P=0.4355). Live cell number and cell viability were confirmed using trypan blue (Fig. 4C). Following 96 hours of recovery in digoxin-free media, cell viability remained high at 103.4±0.73% relative to vehicle control. Although the slight difference at 100 nM digoxin reached formal statistical significance (P=0.0150), significance markers were omitted from the graph because the effect size falls within the inherent margin of error for automated trypan blue counting (∼3-5%) and is considered biologically negligible. As a whole, these results suggest that K562 cell exposure to 100 nM digoxin results in a fully reversible proliferative and metabolic depression.

**Figure 4.**
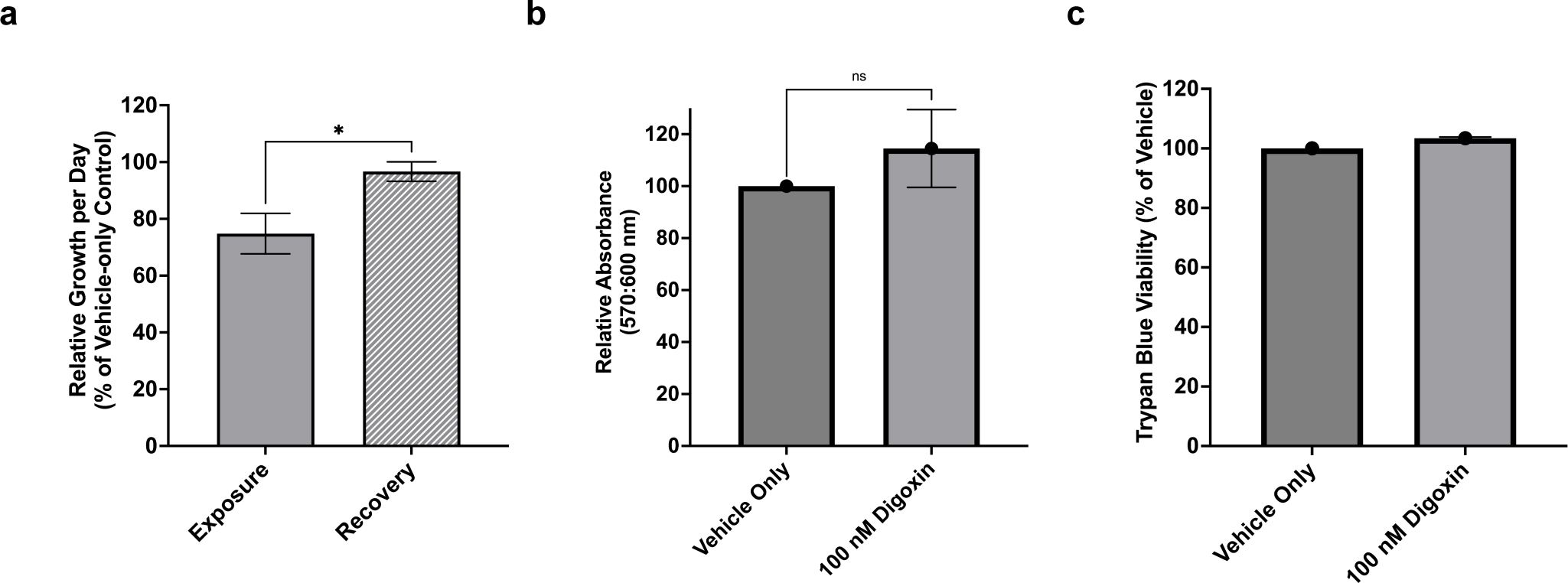
Recovery of K562 growth rate and metabolic activity following digoxin washout. (A) Comparison of K562 cell growth as a percentage of vehicle-only control following 96-hour exposure to 100 nM digoxin, left, or following 96-hour recovery in complete culture media, right. N=3 biological replicates for all groups. (B) Comparison of K562 cell resazurin reduction capacity following 96-hour exposure to digoxin and 96-hour recovery in complete culture media. N=3 biological replicates for all groups. (C) Plot showing viability of cells during recovery, relative to vehicle-only control. N=3 biological replicates for all groups. Statistical significance is represented as P< 0.05 (*), P< 0.01 (**), P< 0.001 (***), P< 0.0001 (****).

## Discussion

In this study, we show that NKA inhibition by digoxin results in dose-dependent attenuation of cancer cell proliferation and reduced metabolic activity at sub-micromolar digoxin concentrations without observable effects on cell viability. This cytostatic growth attenuation and metabolic depression were demonstrated to be fully reversible upon cessation of the pharmacological blockade. By demonstrating that elevated extracellular potassium selectively attenuates digoxin sensitivity in a surmountable manner across both membrane-permeable chloride and membrane-impermeable gluconate counter-anions, we functionally confirm that these growth-inhibitory effects are driven specifically by on-target NKA binding rather than non-specific osmotic shifts. These findings suggest that NKA blockade drives K562 cells towards a temporary oncostatic state, likely via ion dysregulation or non-canonical cell stress signaling.

Overall, this approach presents a novel method and model system for disentangling NKA effects on cancer cell physiology mediated by ion transport and scaffold ligand signaling.

Our observation that 100 nM digoxin slows proliferation without triggering acute cytotoxicity contrasts with some previous reports in leukemia models. For instance, Zhang et al. (2017) reported substantial apoptotic induction when digoxin was combined with PERK inhibitors in K562 cells; however, as a monotherapy at sub-micromolar concentrations (100–500 nM), digoxin induced minimal lethality (<15%), consistent with our finding that single-agent NKA blockade in this concentration window acts primarily via cytostatic attenuation rather than immediate cytotoxic destruction [24]. By directly tracking growth kinetics and metabolic recovery over extended timeframes, our data clarify that K562 cells retain high viability and full proliferative competence upon drug cessation.

### Mechanistic Insights

Maintaining intracellular ion concentrations and transmembrane concentration gradients is energetically costly, but given the essential and ubiquitous functions enabled by a proper intracellular ionic landscape, it is a non-negotiable metabolic investment. Our data clearly showed that NKA blockade attenuates proliferative momentum in K562 cells. In cancer, it stands to reason that NKA function is even more critical, as rapidly dividing cells rely on steep sodium and potassium concentration gradients to facilitate the movement of nutrients and waste across the plasma membrane. Well-characterized functions of ion homeostasis suggest that when ion concentrations become perturbed by NKA pump blockade, the malignancy is starved of nutrients, accumulates toxic metabolic waste, and faces generalized protein instability. Our findings support the premise that one or more of these biophysical constraints force cancer cell populations to limit proliferation rate in order to avoid outright catastrophic collapse of the ionic landscape. When K562 cells are exposed to micromolar concentrations of digoxin, our results suggest that compensatory deceleration is overwhelmed, transitioning the phenotype from reversible cytostasis to metabolic collapse and widespread cytotoxicity.

An alternative mechanism for these observations is through the lens of the signalosome hypothesis, wherein cardiac glycoside binding stimulates non-pumping NKA complexes to activate receptor-bound Src, transactivate EGFR, and engage downstream stress-signaling cascades. In this scenario, increased Src leads to transactivated EGFR and downstream increases in phosphorylation activity. In certain cancer models, excessive mitogenic signaling paradoxically triggers cellular stress and cytostatic arrest. While our data cannot definitively rule out scaffold-receptor induction of cell stress mediated cytostasis, the specific genetic background of K562 cells places substantial constraints on this model. Specifically, K562 cells harbor constitutive BCR-ABL tyrosine kinase activity alongside non-functional TP53 and a homozygous deletion of CDKN2A (p16^INK4a^), meaning that they are largely refractory to canonical kinase-overdrive stress checkpoints. Because kinase-mediated cytostasis relies on intact cell-cycle checkpoints to halt proliferation in response to aberrant kinase cascades, direct biophysical ion disruption seems the more plausible driver of the observed NKA-mediated cytostatic growth attenuation.

### Reversibility, Metabolic Attenuation, and Plasticity

The capacity of K562 cells to fully recover following NKA-mediated cytostasis and metabolic attenuation indicates that sub-micromolar digoxin exposure does not induce irreparable damage or trigger irreversible programmed cell death pathways. Furthermore, these findings support an ion-mediated mechanism, as kinase-driven epigenetic or signaling cascades would likely persist beyond drug washout. While activated Src has previously been linked to suppressed mitochondrial respiration, the immediate restoration of metabolic reducing capacity upon digoxin clearance points toward a biophysical rather than a signaling-locked cause. Plausibly, partial NKA inhibition dissipates the inward sodium gradient required for secondary active transport of glucose and essential amino acids; relieving this pump blockade may promptly re-establish nutrient influx and restore optimal mitochondrial metabolic output. By imposing a reversible bioenergetic choke point, NKA perturbation may effectively constrain the phenotypic plasticity of K562 cells, forcing an adaptive oncostatic pause that could prevent clonal diversification without requiring acute cytotoxic ablation.

### Limitations

Several conceptual and technical limitations of this study warrant consideration. First, while our pharmacological competition assays provide strong functional evidence that NKA inhibition drives the observed cytostatic growth attenuation , we did not directly measure intracellular concentrations of sodium or potassium ions via real-time fluorometry or electrophysiology.

Consequently, our model relies on phenotypic population kinetics and competitive receptor dynamics rather than direct quantification of intracellular ion flux. Second, this investigation was restricted to a single immortalized leukemia cell line. Because K562 cells are rapidly proliferating and metabolically demanding, slower-cycling malignant cell lines or non-proliferative, invasive cancer phenotypes may exhibit altered sensitivity to NKA disruption.

Finally, in cancer types more sensitive to mitogenic stress and harboring intact cell cycle checkpoint machinery, NKA manipulation by cardiac glycosides could plausibly simultaneously engage NKA scaffold receptor-mediated signaling to drive mitogenic stress responses.

### Concluding Remarks and Outlook

From this study, we conclude that digoxin induces a NKA-specific cytostatic growth attenuation and metabolic depression in K562 cells, which are both fully reversible upon insult washout.

These results combined with the unique genetic background of K562 cells demonstrate a novel model system for investigating the NKA, ion homeostasis, and phenotypic plasticity in cancer.

While the specificity of digoxin provides a powerful means for pharmacological dissection of the NKA, notably, our results do not suggest its utility as a cancer therapy. The concentrations of digoxin that induced cytostatic growth attenuation are well above the cardiotoxic serum concentrations observed when digoxin is used to treat heart failure or arrhythmia. However, given the potential mechanism put forth in this study, the identification of a method for inhibiting the NKA without inducing scaffold receptor-mediated signaling may indeed provide better general therapeutic tolerance and prove useful as a cancer therapeutic target. Finally, these findings underscore how systemic electrolyte shifts and altered renal ion handling might serve as crucial physiological indicators to monitor during treatment, while also presenting an exploitable bioenergetic vulnerability where the limits of active NKA compensation can be targeted to constrain tumor growth and phenotypic plasticity.

## Acknowledgements

We thank James Peliska for his insights while interpreting our results. We thank the following University of Mary students for their contributions to this work: Taylor Schmitcke, Jonathan Swartz, Brenden Lucas, Isabelle Parson, Rian Spruenken, Grace Freimuth, Andrijana Fundak, JayCee Frank, Abigail Folk, Elizabeth Folk, Joshua Giacomi, and Blaise Boyle. Research reported in this publication was supported by an Institutional Development Award (IDeA) from the National Institute of General Medical Sciences of the National Institutes of Health under grant number P20GM103442. Additional support was received from the University of Mary.

## Conflicts of Interest

None of the authors have any conflict of interest to disclose.

## Declaration of Generative AI and AI-Assisted Technologies in the Writing Process

During the preparation of this manuscript, the authors used large language model AI tools to assist with language refinement, proofreading, and editorial formatting checks. The authors reviewed and edited all content, and assume full responsibility for the data integrity and conclusions of this publication.

## Notes

### Competing Interest Statement

The authors have declared no competing interest.

### Summary of Updates

Just uploading a word version so that HTML full text can be generated.

## References

1. Greaves, M., & Maley, C. C. (2012). Clonal evolution in cancer. Nature, 481(7381), 306–313. 10.1038/nature10762

2. Gillies, R. J., Verduzco, D., & Gatenby, R. A. (2012). Evolutionary dynamics of carcinogenesis and why targeted therapy does not work. Nature Reviews Cancer, 12(7), 487–493. 10.1038/nrc3298

3. Hanahan, D. (2026). Hallmarks of cancer—Then and now, and beyond. Cell, 189(3), 1–18. 10.1016/j.cell.2025.12.049

4. Prevarskaya, N., Skryma, R., & Shuba, Y. (2018). Ion channels in cancer: Are cancer hallmarks oncochannelopathies? Physiological Reviews, 98(2), 559–621. 10.1152/physrev.00044.2016

5. Ruffinatti, F. A., Scarpellino, G., Chinigò, G., Visentin, L., & Munaron, L. (2023). The emerging concept of transportome: state of the art. Physiology, 38(6), 285–302. 10.1152/physiol.00010.2023

6. Rolfe, D. F. S., & Brown, G. C. (1997). Cellular energy utilization and molecular origin of standard metabolic rate in mammals. Physiological Reviews, 77(3), 731–758. 10.1152/physrev.1997.77.3.731

7. Yang, M., & Brackenbury, W. J. (2013). Membrane potential regulates cell cycle progression in human cells. Frontiers in Physiology, 4, 185. 10.3389/fphys.2013.00185

8. Blackiston, D. J., McLaughlin, K. A., & Levin, M. (2009). Bioelectric controls of cell proliferation: ion channels, membrane voltage and the cell cycle. Cell Cycle, 8(21), 3527–3536. 10.4161/cc.8.21.9888

9. Lang, F., Busch, G. L., Ritter, M., et al. (1998). Functional significance of cell volume regulatory mechanisms. Physiological Reviews, 78(1), 247–306. 10.1152/physrev.1998.78.1.247

10. Hoffmann, E. K., Lambert, I. H., & Pedersen, S. F. (2009). Physiology of cell volume regulation in vertebrates. Physiological Reviews, 89(1), 193–277. 10.1152/physrev.00037.2007

11. Prassas, I., & Diamandis, E. P. (2008). Novel therapeutic applications of cardiac glycosides. Nature Reviews Drug Discovery, 7(11), 926–935. 10.1038/nrd2682

12. Babula, P., Masarik, M., Adam, V., et al. (2013). From Na+/K+-ATPase and cardiac glycosides to cytotoxicity and cancer treatment. Anti-Cancer Agents in Medicinal Chemistry, 13(7), 1069–1087. 10.2174/18715206113139990304

13. Li, Z., & Xie, Z. (2009). The Na/K-ATPase/Src complex and cardiotonic steroid-activated protein kinase cascades. Pflügers Archiv - European Journal of Physiology, 457(3), 635–644. 10.1007/s00424-008-0470-0

14. Cui, X., & Xie, Z. (2017). Protein interaction and Na/K-ATPase-mediated signal transduction. Molecules, 22(6), 990. 10.3390/molecules22060990

15. Haas, M., Askari, A., & Xie, Z. (2000). Involvement of Src and epidermal growth factor receptor in the signal-transducing function of Na+/K+-ATPase. Journal of Biological Chemistry, 275(36), 27832–27837. 10.1074/jbc.M002951200

16. Tian, J., Cai, T., Yuan, Z., et al. (2006). Binding of Src to Na+/K+-ATPase forms a functional signaling complex. Molecular Biology of the Cell, 17(1), 317–326. 10.1091/mbc.e05-08-0735

17. Liang, M., Tian, J., Liu, L., et al. (2007). Identification of a pool of non-pumping Na/K-ATPase. Journal of Biological Chemistry, 282(14), 10585–10593. 10.1074/jbc.M609181200

18. Bartlett, D. E., et al. (2018). The role of Na/K-ATPase signaling in oxidative stress related to aging: Implications in obesity and cardiovascular disease. IJMS, 19(7), 2139. 10.3390/ijms19072139

19. Kometiani, P., Liu, L., & Askari, A. (2005). Digitalis-induced signaling by Na+/K+-ATPase in human breast cancer cells. Molecular Pharmacology, 67(3), 929–936. 10.1124/mol.104.007302

20. Geng, X., Wang, F., Tian, D., Huang, L., Streator, E., Zhu, J., Kurihara, H., He, R., Yao, X., Zhang, Y., & Tang, J. (2020). Cardiac glycosides inhibit cancer through Na/K-ATPase-dependent cell death induction. Biochemical Pharmacology, 182, 114226. 10.1016/j.bcp.2020.114226

21. Wang, Y., Hou, Y., Hou, L., Wang, W., Li, K., Zhang, Z., Du, B., & Kong, D. (2021). Digoxin exerts anticancer activity on human nonsmall cell lung cancer cells by blocking PI3K/Akt pathway. Bioscience Reports, 41(10), BSR20211056. 10.1042/BSR20211056

22. Prassas, I., Karagiannis, G. S., Batruch, I., Dimitromanolakis, A., Datti, A., & Diamandis, E. P. (2011). Digitoxin-induced cytotoxicity in cancer cells is mediated through distinct kinase and interferon signaling networks. Molecular Cancer Therapeutics, 10(11), 2083–2093. 10.1158/1535-7163.MCT-11-0421

23. Lei, Y., Gan, H., Huang, Y., Chen, Y., Chen, L., Shan, A., Zhao, H., Wu, M., Li, X., Ma, Q., Wang, J., Zhang, E., Zhang, J., Li, Y., Xue, F., & Deng, L. (2020). Digitoxin inhibits proliferation of multidrug-resistant HepG2 cells through G_2_/M cell cycle arrest and apoptosis. Oncology Letters, 20(4), 71. 10.3892/ol.2020.11932

24. Zhang, X.-H., Wang, X.-Y., Zhou, Z.-W., Bai, H., Shi, L., Yang, Y.-X., Zhou, S.-F., & Zhang, X.-C. (2017). The combination of digoxin and GSK2606414 exerts synergistic anticancer activity against leukemia in vitro and in vivo. BioFactors, 43(6), 812–820. 10.1002/biof.1380

25. Amarante-Mendes, G. P., Kim, C. N., Liu, L., et al. (1998). Bcr-Abl exerts its antiapoptotic effect against diverse apoptotic stimuli through blockage of mitochondrial release of cytochrome C and activation of caspase-3. Blood, 91(5), 1700–1705. 10.1182/blood.V91.5.1700

26. Deininger, M. W., Goldman, J. M., & Melo, J. V. (2000). The molecular biology of chronic myeloid leukemia. Blood, 96(10), 3343–3356. 10.1182/blood.V96.10.3343

27. Lozzio, C. B., & Lozzio, B. B. (1975). Human multipotential leukemia cell line (K-562) with Philadelphia chromosome. Blood, 45(3), 321–334. 10.1182/blood.V45.3.321.321

28. Law, J. C., Ritke, M. K., Yalowich, J. C., Leder, G. H., & Ferrell, R. E. (1993). Mutational inactivation of the p53 gene in the human erythroid leukemic K562 cell line. Leukemia Research, 17(12), 1045–1050. 10.1016/0145-2126(93)90161-d

29. Sill, H., Goldman, J. M., & Cross, N. C. (1995). Homozygous deletions of the p16 tumorsuppressor gene are associated with lymphoid transformation of chronic myeloid leukemia. Blood, 85(8), 2013–2016. 10.1182/blood.V85.8.2013.bloodjournal8582013

30. Ogawa, H., Shinoda, T., Cornelius, F., & Toyoshima, C. (2009). Crystal structure of the sodium-potassium pump Na+/K+-ATPase with bound potassium and ouabain. Proceedings of the National Academy of Sciences, 106(33), 13742–13747. 10.1073/pnas.0907054106

31. Gardner, J. D., & Conlon, T. P. (1972). Effects of sodium and potassium on ouabain and digoxin binding to human red blood cells. Journal of General Physiology, 60(5), 609–629. 10.1085/jgp.60.5.609

32. Rampersad, S. N. (2012). Multiple applications of Alamar Blue as an indicator of metabolic function and cellular health in cell viability/metabolic in vitro assays. Sensors, 12(9), 12347–12360. 10.3390/s120912347

33. Valente, L. C., Paes, M. C., & Maya-Monteiro, C. M. (2017). Ouabain-mediated Na+/K+-ATPase inhibition induces metabolic shifts and mitochondrial dysfunction in cancer cells. Biochimie, 137, 169–178. 10.1016/j.biochi.2017.04.012

